# Is there a negative trend in abundance of the weedy seadragon (*Phyllopteryx taeniolatus*) discernible in survey data summarised in Edgar et al. (2023)?

**DOI:** 10.1101/2024.07.21.604472

**Authors:** Erik Schlögl, David J. Booth

## Abstract

Decadal changes in abundance of marine organisms along the Australian coastline were reported in a recent *Nature* paper by Edgar et al. (2023). We reexamine the data used in that paper, and show that the conclusion of a 59% decline in weedy seadragons (*Phyllopteryx taeniolatus*) cannot be justified based on these data. The data are too sparse, especially for the final survey years of 2020 and 2021, and it is this sparsity in combination with particular modelling choices made by Edgar et al. (2023) that drives their results. While not specifically considered here, it is likely that these issues also affect the results on numerous other species reported in Edgar et al. (2023).

## 1 Introduction

In their paper, “Continent–wide declines in shallow reef life over a decade of ocean warming”, Edgar et al. (2023) state that in their dataset encompassing Australia’s nearshore seas, “Populations of 57% of reef species decreased in the decade to 2021,” and, “Populations of 28% of [marine] species declined by more than 30% between 2011 and 2021 (9 corals, 36 invertebrates, 24 macroalgae and 227 vertebrates), thus passing the threshold that qualifies species as threatened for the [International Union for the Conservation of Nature (IUCN)] Red List of Threatened Species when generation length is unknown.” As a notable example, they mention “the weedy seadragon (*Phyllopteryx taeniolatus*), an iconic southern Australian endemic fish that significantly declined by 59% from 2011 to 2021”. This species was most recently assessed for *The IUCN Red List of Threatened Species* in 2016 and is listed as “Least Concern”.

The raw data for *P. taeniolatus* used by Edgar et al. (2023) contains 53 rows, i.e. 53 distinct combinations of survey site and survey method. There is a column for each year from 1992 to 2021, although pre-2008 data are not referenced in the paper (in fact, the statement quoted above refers to the eleven years from 2011 to 2021). In a year, on average 17.47 of the 53 distinct site/method combinations were completed (i.e. had actual data). Thus, the data are very sparse overall. For 2021, this is especially so, with only 7 rows containing actual data, i.e., 7 sites were surveyed. Furthermore, at all 7 of these sites, no *P. taeniolatus* were counted in 2020 or 2021. However, even if we assume that there was no decline in abundance of *P. taeniolatus*, this is not an unlikely outcome, because at those seven sites, in the majority of past years (on average more than 80%) where those sites were surveyed, the count of *P. taeniolatus* was zero. The sparsity of the data and the substantial year–on–year variation of the observed^1^ abundance are illustrated in Figure 1.

**Figure 1:**
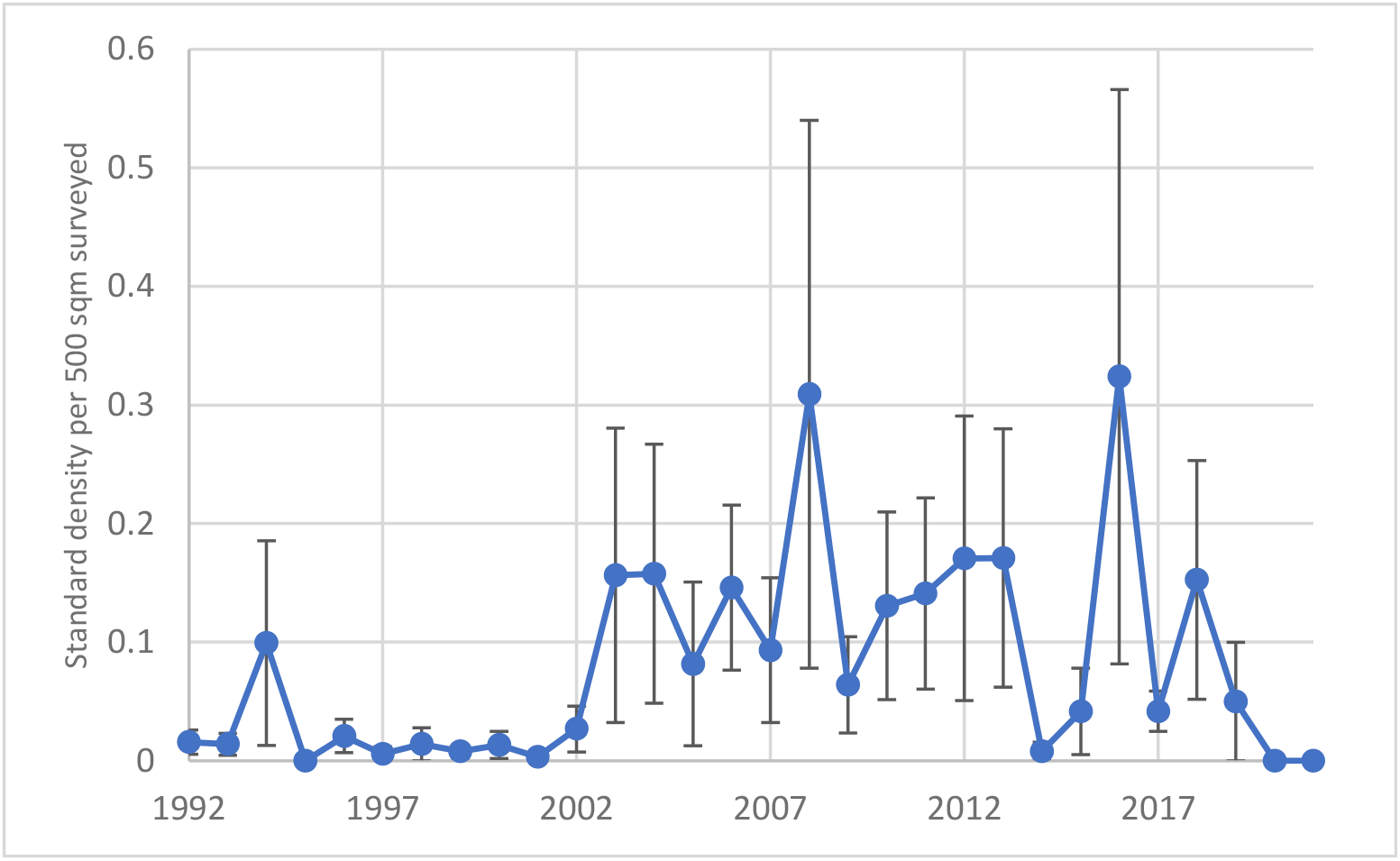
Standard density of *P. taeniolatus* per 500m^2^ surveyed (i.e., raw data as used by Edgar et al. (2023)), averaged across all survey sites actually surveyed in each given year. Error bars represent *±*2 standard deviations of the yearly sample average.

The sparsity of the data (and, arguably, visual inspection of Figure 1) suggests that concluding that this species declined by 59% between 2011 and 2021 (as Edgar et al. (2023) claim) is not supported by the data, but rather may be an artefact of their data aggregation, choice of time window, interpolation and extrapolation. This note seeks to explore this issue.

## 2 Statistical significance of downward population trend

Based on the data used by Edgar et al. (2023), no rigorous statistical test would lead to rejection of a null hypothesis that *P. taeniolatus* declined by less 30% (the IUCN threshold quoted in the paper) between 2011 and 2021. It is worth noting that based on the information provided with the paper, it also does not seem that Edgar et al. (2023) tested that particular hypothesis. We have attempted to reproduce the statistical test for this species that they did conduct, as per the R code provided by the authors on Github (we recoded this in Python). This is a Spearman’s rank correlation test^2^ conducted on data that were not interpolated or extrapolated, but aggregated so that there is one data point per year. This aggregation is still problematic, but (for the sake of argument) accepting the approach of Edgar et al. (2023), we find no statistically significant decline in *P. taeniolatus* based on the data from 2011 to 2021. The *p*-value is 0.23 using permutation rather than asymptotic methods, and 0.47 using an asymptotic method (the former is more appropriate due to the small number of data points). However, based on the information provided on Github, there seems to be a discrepancy between the statistical test that the authors ran and how they reported this in the paper: The information on Github suggests that they ran the test based on data from 2008 to 2021, on which we also obtain a significant decline, with a (permutation method) *p*-value of 0.035. For any “decadal” window, however, this significance goes away: 2008 to 2018 gives a *p*-value of 0.42, 2009 to 2019 a *p*-value of 0.15, and 2010 to 2020 a *p*-value of 0.17. Thus, using the statistical test suggested by Edgar et al. (2023), one cannot exclude the substantial possibility that any observed downward trend in abundance of *P. taeniolatus* is simply due to random noise, rather than an actual decline.

## 3 Robustness of the population trend estimate

Based on Section 3.2.4 “Dealing with uncertainty” of the Red List Guidelines (see IUCN Standards and Petitions Committee (2022)), statistical significance of a downward trend may be too high a bar for a threatened species listing. This section states,

> *“It is recommended that assessors should adopt a precautionary but realistic attitude, and to resist an evidentiary attitude to uncertainty when applying the criteria (i.e*., *have low risk tolerance). This may be achieved by using plausible lower and upper bounds, rather than only the best estimates, in determining the quantities used in the criteria. In cases where a statistical method is used to estimate a quantity, a 90% confidence interval or 90% credible interval may be used to set the plausible range of values. It is recommended that ‘worst case scenario’ reasoning be avoided because this may lead to unrealistically precautionary listings. All attitudes should be explicitly documented.”*

Thus, the Guidelines suggest using a trend value such that we can be 90% sure (based on the data) that the true trend is better (e.g. less downward sloping) than this. However, this reinforces the point that the data used in Edgar et al. (2023) are insufficient (i.e., too sparse) to draw any meaningful conclusions about the population trend of for example *P. taeniolatus* one way or the other — a correctly determined confidence interval at the 90% level based on the data used in the paper will far too wide to be meaningful, and indeed Edgar et al. (2023) do not provide such confidence intervals.

Instead, Edgar et al. (2023) provide what they appear to consider a best estimate of the population trend, and it is to this estimate that they are referring in their statement, “the weedy seadragon (*Phyllopteryx taeniolatus*), an iconic southern Australian endemic fish that significantly declined by 59% from 2011 to 2021”. As noted above, “significantly” in this context cannot be interpreted as statistical significance, so the question to now consider is whether this is indeed a reasonably robust best estimate of the population decline, or rather an artefact of their modelling choices.

One can show that the latter is the case through a relatively simple exercise accepting (for the sake of argument) all of their modelling choices and varying the data time window for the trend estimation. Thus, we follow Edgar et al. (2023) and first interpolate and extrapolate the data they have used in the same manner as they do, i.e., for years where there are no data available for a particular survey site, linear interpolation between the closest preceding and following years for which data is available is performed. Where there is no preceding year with data available, the value is set equal to value for the closest following year with data, and where there is no following year with data available, the value is set equal to value for the closest preceding year with data. After rounding latitude and longitude of each survey site to the closest full degree, sites with the same rounded latitude and longitude are grouped together, and the average value for each year is calculated for each such group. Subsequently, these group averages for each year are averaged across all groups, to yield a series of single values per year. After taking the natural logarithm of these values, we find the best–fitting straight line (in a least–squares sense), i.e., we find the slope of the linear regression of the logarithmic observation values on the observation year. Reading off the value *y*_start_ on the regression line at the starting year and the value *y*_end_ on the regression line at the end year, the estimated relative change in abundance is given by

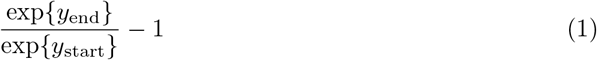

Due to the sparsity of the data, this estimate is very sensitive to the data window on which the above procedure is applied. Consider Table 1, which gives the abundance trend estimates with different data window length, all ending 2021. If one uses the data from 2015 to 2021, there is an estimated decline of 50.7% between the starting year and the end year, while between 2011 and 2021 the estimated decline is actually less, 34.3%.^3^ Note that we are unable to reproduce the estimate of a 59% decline between 2011 and 2021 stated by Edgar et al. (2023).

**Table 1:**
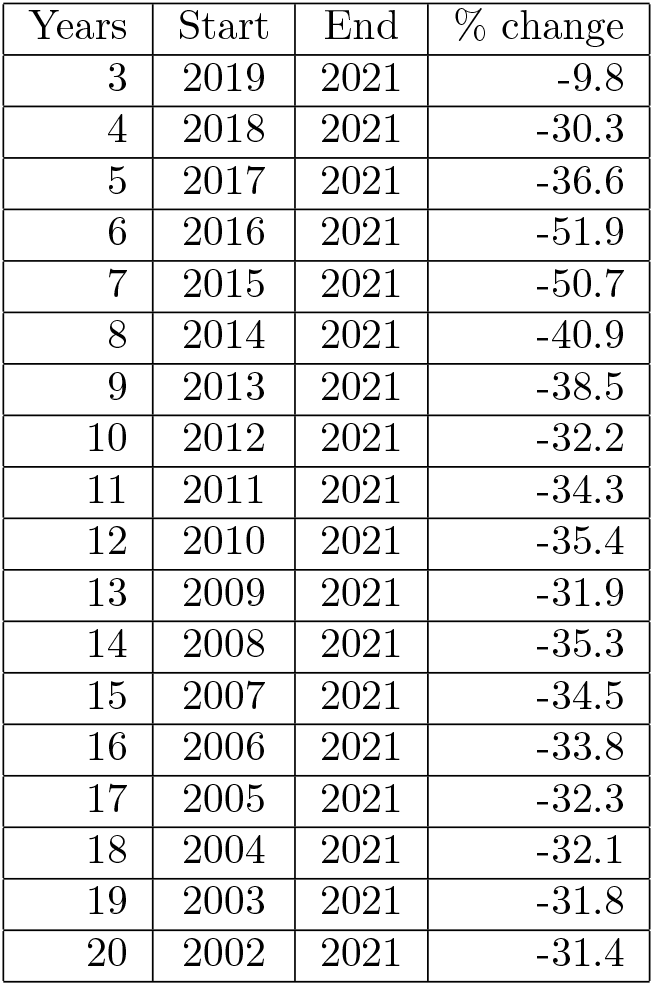
Abundance trend estimates using the method described in Edgar et al. (2023) with different data window length, all ending 2021.

| Years | Start | End | % change |
| --- | --- | --- | --- |
| 3 | 2019 | 2021 | -9.8 |
| 4 | 2018 | 2021 | -30.3 |
| 5 | 2017 | 2021 | -36.6 |
| 6 | 2016 | 2021 | -51.9 |
| 7 | 2015 | 2021 | -50.7 |
| 8 | 2014 | 2021 | -40.9 |
| 9 | 2013 | 2021 | -38.5 |
| 10 | 2012 | 2021 | -32.2 |
| 11 | 2011 | 2021 | -34.3 |
| 12 | 2010 | 2021 | -35.4 |
| 13 | 2009 | 2021 | -31.9 |
| 14 | 2008 | 2021 | -35.3 |
| 15 | 2007 | 2021 | -34.5 |
| 16 | 2006 | 2021 | -33.8 |
| 17 | 2005 | 2021 | -32.3 |
| 18 | 2004 | 2021 | -32.1 |
| 19 | 2003 | 2021 | -31.8 |
| 20 | 2002 | 2021 | -31.4 |

Furthermore, the approach to missing data chosen by Edgar et al. (2023), i.e. filling in missing data by interpolation and extrapolation, will tend to reinforce spurious trend estimates. This is particularly true of extrapolation. The observation data of *P. taeniolatus* used by Edgar et al. (2023) is especially sparse in the final two years (i.e., 2020 and 2021), so to mitigate the impact of extrapolation (somewhat), one could consider only the data up to 2019, which gives an estimated abundance reduction of 17.8% from the baseline year of 2008 (and only 11.7% when considering the “decadal” period starting 2009). However, from Table 2 it is clear that trend estimates are highly variable depending on the choice of data window. This is true even when comparing data windows that largely overlap and only differ by a single year. Thus, the data used by Edgar et al. (2023) are insufficient to obtain any informative estimate of a trend in the abundance of *P. taeniolatus* and thus do not support their conclusion of a substantial decline.

**Table 2:** Abundance trend estimates using the method described in Edgar et al. (2023) with different data window length, ending in different years.

| Start | % change to 2017 | % change to 2018 | % change to 2019 | % change to 2020 |
| --- | --- | --- | --- | --- |
| 2015 | -11.8 | -21.1 | -38.2 | -47.1 |
| 2014 | 20.2 | 3.7 | -20.4 | -34.1 |
| 2013 | 18.0 | 5.0 | -17.2 | -31.1 |
| 2012 | 27.4 | 14.8 | -8.1 | -23.4 |
| 2011 | 13.7 | 5.4 | -12.9 | -26.2 |
| 2010 | 5.9 | -0.2 | -15.8 | -27.8 |
| 2009 | 9.7 | 3.7 | -11.7 | -23.9 |
| 2008 | -1.1 | -5.2 | -17.8 | -28.3 |
| 2007 | -1.9 | -5.6 | -17.5 | -27.5 |

## 4 Conclusion

In summary:

- Based on the data used by Edgar et al. (2023), one cannot conclude that the population of *P. taeniolatus* has declined in a way that should trigger an IUCN classification of “threatened”. The data are too sparse.
- The statement about the magnitude of decline in this species in Edgar et al. (2023) is an artefact of the way they treat the data.
- A statistically significant decline of any magnitude of this species over a “decadal” period is not supported by the data, even if using their (flawed) methodology.

It seems that at least some of the issues identified with the statistical analysis of the *P. taeniolatus* carry over to numerous other species in the dataset used by Edgar et al. (2023): Specifically, for numerous other species, the data are also very sparse, so any abundance trend estimates based on such data will be similarly lacking in robustness. Using the methodology of Edgar et al. (2023), random observation error alone would result in negative trend estimates for around 50% of species considered *even if there is no actual change in abundance of any species*, and looking at the entirety of species considered in Edgar et al. (2023), they note, “Populations of 57% of reef species decreased in the decade to 2021, a percentage that was consistent whether all species (617 out of 1,057, 58%) or only species with significant trends (*P <* 0.05; 97 out of 172, 57%) were considered.” Thus, while it is reasonable to expect some of the qualitative results which they obtain with respect to changes in species abundance as a consequence of ocean warming, from a quantitative/statistical perspective the data do not support their conclusions, in particular (but not limited to) with respect to the 59% decline of *P. taeniolatus* highlighted in the paper.

The fact that the analysis documented in this note demonstrates that the population decline of *P. taeniolatus* reported in Edgar et al. (2023) is not scientifically defensible is of special concern, as this purported decline was widely reported in the media.^4^ Errors of fact in research on climate change impacts can contribute to public scepticism with regard to this (scientifically well established) phenomenon. One course of action would be for Edgar et al. (2023) to retract their conclusions as appropriate.

## Code and data availability

The code and data used to prepare this note are available at https://github.com/eschlogl/WeedySeadragon. The code is in Python, in the form of a Jupyter notebook `WeedySeadragon.ipynb`, also provided in read-only form as an HTML file `WeedySeadragon.html`.

## Footnotes

1 Note that *P. taeniolatus* sexually mature at 28 months and tagging studies have shown a longevity of up to six years (see Sanchez-Camara et al. (2005)). Thus, the abundance spikes in 2008 and 2016 evident in Figure 1 are likely due to noise in the survey data, rather than actual changes in abundance.

2 Recall that the Spearman’s rank correlation test only can make a statement as to whether there was a statistically significant decline or increase, but is silent about the magnitude of such a decline or increase.

3 Note that the period from 2015 to 2021 is included in the period from 2011 to 2021, and therefore the only way we can have a 50% decline in the shorter period and a 34% decline in the longer period is if there was a substantial abundance increase between 2011 and 2015, which would mean that if one accepts the approach of Edgar et al. (2023), one would conclude that there are cyclical abundance variations, (or at least variations not related to ocean warming) on the order of 20%. However, we consider it most likely both the increase and the decrease are primarily due to observation noise and sparse data, exacerbated by an inappropriate methodology.

4 See e.g. Kilvert (2023), Alberts (2023) or the lead author’s own article in *The Conversation*, Edgar (2023). Readfearn (2023) also used an image of *P. taeniolatus* as the main illustration accompanying an article on the findings of Edgar et al. (2023).

